# The Hsp90 chaperone system from the African trypanosome, *Trypanosoma brucei*

**DOI:** 10.1101/2021.03.15.435448

**Authors:** Miebaka Jamabo, Stephen J. Bentley, Paula Macucule-Tinga, Adrienne L. Edkins, Aileen Boshoff

## Abstract

African Trypanosomiasis is a neglected tropical disease caused by *Trypanosoma brucei* (*T. brucei*) and is spread by the tsetse fly in sub-Saharan Africa. The disease is fatal if left untreated and the currently approved drugs for treatment are toxic and difficult to administer. The trypanosome must survive in the insect vector and its mammalian host, and to adapt to these different conditions, the parasite relies on molecular chaperones called heat shock proteins. Heat shock proteins mediate the folding of newly synthesized proteins as well as prevent misfolding of proteins under normal conditions and during stressful conditions. Heat shock protein 90 (Hsp90) is one of the major molecular chaperones of the stress response at the cellular level. It functions with other chaperones and co-chaperones and inhibition of its interactions is being explored as a potential therapeutic target for numerous diseases. This study provides an in-silico overview of Hsp90 and its co-chaperones in both *T. brucei brucei and T. brucei gambiense* in relation to human and other kinetoplastid parasites. The evolutionary, functional, and structural analyses of Hsp90 were also shown. The updated information on Hsp90 and its co-chaperones from recently published proteomics on *T. brucei* was examined for the different life cycle stages and subcellular localisations. The results show a difference between *T. b. brucei and T. b. gambiense* with *T. b. brucei* encoding 12 putative *Hsp90* genes, 10 of which are cytosolic and located on a single chromosome while *T. gambiense* encodes 5 *Hsp90* genes, 3 of which are located in the cytosol. Eight putative co-chaperones were identified in this study, 6 TPR-containing and 2 non-TPR-containing co-chaperones. This study provides an updated context for studying the biology of the African trypanosome and evaluating Hsp90 and its interactions as potential drug targets.

## Introduction

*Trypanosoma brucei* (*T. brucei*), is an extracellular blood- and tissue-borne protozoan parasite transmitted by tsetse fly vectors, which causes devastating diseases in humans, wild animals and domesticated livestock (1). Human African trypanosomiasis (HAT, although known as African sleeping sickness), is a potentially fatal tropical disease found in remote rural regions of sub-Saharan Africa and often coincides with insubstantial health care systems (2). HAT is caused by two subspecies of *T. brucei*; The chronic form of the disease, which is endemic to Central and Western Africa, is caused by *Trypanosoma brucei* (*T. b.*) *gambiense*, and the acute zoonotic form, which is endemic to Eastern and Southern Africa, is caused by *T. b. rhodesiense* (3, 4). The livestock disease Nagana, caused by *T. b. brucei* also has a crippling effect on the socioeconomic development within sub-Saharan Africa (5, 6). Despite the decreasing number of HAT cases, there is still a desperate need for the development of new and more effective drugs due to the difficult administration and toxicity of the current treatments, lack of a vaccine and increasing parasite resistance (7). Molecular chaperones have been identified as an attractive target for drug development against protozoan parasites as this protein family plays essential roles in stress-induced stage differentiation and are vital for disease progression and transmission (8–10).

The 90-kDa heat shock protein (Hsp90) family contains essential, highly conserved and abundant molecular chaperones (11–13) that facilitate the proper folding and maturation of a large but specific group of substrates called client proteins (14–16). More than 400 client proteins have been identified to date (listed at http://www.picard.ch/), with many of them being implicated in protein folding and degradation, signalling pathways, cellular trafficking, cell cycle regulation, differentiation, and others (17–19). In eukaryotes, the Hsp90 family is normally comprised of four isoforms that are located in various cellular compartments. Two Hsp90 (the stress-inducible form Hsp90α/HSPC2 and the constitutive form Hsp90β/HSPC3) isoforms are located in the cytosol and in the nucleus (20–22); GRP94/HSPC4 is present in the endoplasmic reticulum (ER) (20,21,23) and TRAP1/HSPC5 is found in the mitochondrial matrix (24). Some intracellular Hsp90 isoforms are exported and function in the extracellular environment to regulate the immune response, cell migration and invasion (25–28).

Structurally, Hsp90 is a flexible dimeric protein with each monomer containing three domains: an N-terminal nucleotide-binding domain (NBD); a middle client protein-binding domain (MD); and a C-terminal dimerization domain (DD) (29–31). To perform its molecular chaperone function, Hsp90 is dependent on ATP hydrolysis, and a battery of accessory proteins termed co-chaperones, which assist in the recruitment of client proteins and the regulation of the Hsp90 reaction cycle (32, 33). The cytosolic Hsp90 isoforms contain a conserved C-terminal MEEVD motif which acts as a docking site for interaction with co-chaperones that possess the tetratricopeptide repeat-(TPR) domain (34, 35). Other Hsp90 co-chaperones interact with the molecular chaperone through its NBD or M domain (33). So far, more than 30 co-chaperones have been identified in the mammalian Hsp90 chaperone system. However, the composition of the Hsp90 chaperone system appears to vary across organisms indicating that the function of some co-chaperones may be restricted to specific subsets of client proteins, be required for client protein activation in a species-dependent manner, or be redundant with other co-chaperones (36). Hsp90 is also subject to post-translational modifications, including s-nitroslyation, phosphorylation and acetylation, which may influence its activity, cellular localization or its interaction with co-chaperones, nucleotides or client proteins (37–40). Some Hsp90 isoforms are essential for viability, and maintenance of client proteins that are dependent on the chaperone (41), making it an attractive drug target for diseases including infectious diseases. Several Hsp90 inhibitors, which have been well studied in the laboratory and clinic for antitumor indications (42, 43), were also shown to arrest the growth of several kinetoplastids *in vitro* and have activity against *Trypanosoma evansi* and *T. brucei* in mice (44–47). Thus, the repurposing of Hsp90 inhibitors designed for cancer treatment is one strategy to evaluate new and effective anti-trypanosomal agents (48).

Post-genomic analysis of the molecular chaperone complements in kinetoplastid parasites have revealed unprecedented expansion and diversification, highlighting the importance of these protein families in the biology of these organisms (8,9,49–51). In *Trypanosoma* and *Leishmania*, the Hsp90 (Hsp83) machinery has a pivotal role in environmental sensing and life cycle control (44,52,53). Several reviews and updated *in silico* analyses of the Hsp90/HSPC family and Hsp90 heterocomplexes in the annotated genome sequences of intracellular kinetoplastid parasites have been conducted (8,50,51,54,55). However, this has not been the case for the extracellular parasite, *T. brucei*.

*T. brucei* exhibits a digenetic lifestyle, and therefore must adapt to fluctuating environmental conditions, such as change in temperature, pH, nutrients and the pressure from the immune system, as it transitions from the gut of the tsetse fly to the body fluids of its mammalian host (54, 56). A distinct molecular trait of trypanosomes is their dependence on polycistronic transcription akin to prokaryotes. Trypanosomal mRNAs are mainly generated through trans-splicing and there is a dependence on post-transcriptional mechanisms for gene regulation (57). However, correlation studies comparing the previously reported RNA-seq data of transcript abundance and proteomic data from the procyclic form (PF) and bloodstream form (BSF) of the parasite show that the differences observed between the PF and BSF are two-fold greater at the proteomic level when compared to the transcriptomic level (58, 59). Given the complexities of transcription, its incomplete representation of the life cycle stages of the parasite as well as its lack of control, trypanosome research has largely shifted to rely on proteomic data (60). Numerous proteomic studies have been conducted in the parasite which have compared protein expression at the different life cycle stages (58,59,61), in the mitochondrion (62), mitochondrial importome (63), respiratome (64), mitochondrial membranes (outer, intermembrane space, inner and matrix) (65), nucleus (60), nuclear pore (66), glycosomes (67, 68), flagellum (69, 70) and cell surface (71).

*T. brucei* and the related kinetoplastids rely on post-translational modifications (PTMs) to increase their proteome diversity and complexity (72). Advanced studies in trypanosomes phosphoproteome and acetylome (73–76) indicates phosphorylation and acetylation as the most predominant modifications in *T. brucei* proteins. Both PTMs are well known for impacting Hsp90 intracellular localization as well as their ability to bind co-chaperones, nucleotides, clients (72, 74) and even inhibitors (77). However, the PTMs regulatory dynamic in the organellar TRAP-1 and GRP94 are yet to be elucidated for a global understanding of this critical chaperone activity regulator system.

This paper aimed to provide a comprehensive depiction of the *T. brucei* Hsp90 chaperone system based on structural, functional, and evolutionary analyses. *In silico* tools were used to evaluate the domain conservation, predicted subcellular localisations, syntenic and phylogenetic analysis of the Hsp90 chaperone system in *T. brucei* with respect to both *T. b. brucei* and *T. b. gambiense*. The Hsp90 chaperone system was also comparatively analysed in relation to those found in selected kinetoplastid parasites and *Homo sapiens*. The proteomic findings on Hsp90 and its co-chaperones from the numerous published proteomic data on *T. brucei* are presented, and we provide updated insights on the adaptability of the parasite from its stage-specific expressed proteins and overall provide a context for identifying new and potential drug targets for HAT.

## Materials and Methods

### Database mining, sequence analyses and the determination of the kinetoplastid and human orthologues

A BLASTP search using the amino acid sequences of Hsp90 isoforms from the *T. b. brucei* obtained from previous *in silico* study (49), and the human HSPC2, HSP90AB1/HSPC3, HSP90B1/HSPC4 and HSPC5 isoforms were used as queries on the TriTrypDB (version 46) database (https://tritrypdb.org/tritrypdb/) (78) and were analyzed in order to determine the Hsp90 complement encoded on the *T. b. gambiense* genome, as well as identify new *T. b. brucei* Hsp90/HSPC protein members. The e-value was set at a stringent level of e^-10^ to identify potential Hsp90/HSPC-related sequences for further analysis. Additionally, a keyword search was performed to scan the genome of *T. b. gambiense* for Hsp90/HSPC genes on the TriTrypDB using the search terms: “Hsp90”, “Hsp83”, “heat shock protein” and “molecular chaperone”. The retrieved amino acid sequences from the various keyword searches were screened using SMART 7 (Simple Modular Architecture Research Tool; http://smart.embl-heidelberg.de/) (79) and PROSITE (http://prosite.expasy.org/) (80) for domains annotated by the online servers as “Hsp90”.

For identification of *T. brucei* orthologs of selected cytosolic Hsp90 co-chaperones, the protein sequences of the human co-chaperones were used as queries in a BLASTP search on the TriTrypDB. Reciprocal BLASTP was conducted to determine if the identified putative *T. brucei* co-chaperone had the closest match to the desired human co-chaperone. The putative amino acid sequences of the co-chaperones from both *T. brucei* subspecies were used as queries in a BLASTP search on the National Centre for Biotechnology Information (NCBI) website (www.ncbi.nlm.nih.gov), using the default parameters. If the most similar orthologue in the *T. brucei* subspecies was identical to the Hsp90 co-chaperone sequence used as first query, the sequence of the second query was selected as an orthologue. Reciprocal BLASTP was also conducted for the identification of human and selected kinetoplastid orthologues of the putative Hsp90/HSPC proteins from both *T. brucei* subspecies.

### Phylogenetic and conserved syntenic analysis

A phylogenetic tree was constructed to analyse the phylogenetic relationship of the Hsp90/HSPC complements in both *T. brucei* subspecies. The full-length amino acid sequences for the Hsp90/HSPC family in the selected kinetoplastid parasites were obtained from TriTryDB (78), and the human protein sequences were obtained from the NCBI website (www.ncbi.nlm.nih.gov). Partial amino acid sequences were omitted from the analysis. Gene ID numbers for the Hsp90/HSPC sequences used in this study are provided in Table 1. Multiple sequence alignments were performed using the inbuilt ClustalW program (81) with default parameters in MEGA-X (82), and is provided in the supplementary data, Fig S1. Maximum likelihood (ML) was utilized to find the best model of evolution and was selected by the Bayesian Information Criterion (BIC) implemented in MEGA-X. The amino acid-based Hsp90/HSPC ML phylogeny was reconstructed using the JTT (Jones-Taylor-Thornton) model matrix (83) with gamma distribution shape parameter (G). The ML phylogenetic tree was constructed using MEGA-X (82). The accuracy of the reconstructed tree was assessed using a bootstrap test using 1000 replicates with a pairwise gap deletion mode. The phylogenetic tree for the Hsp90s was unrooted.

**Table 1.** The Hsp90/HSPC proteins from *Trypanosoma brucei* with putative orthologues in *T. cruzi*, *L. major*, *C. fasciculata*, *B. saltans* and *H. sapiens*.

|  | *H. sapiens* | *T. brucei* | *T. cruzi*<sup>c</sup> | *L. major* | *C. fasciculata* | *B. saltans* |  |  |
| --- | --- | --- | --- | --- | --- | --- | --- | --- |
| Name<sup>a</sup> | Gene ID<sup>b</sup> | Gene ID<sup>b</sup> | Gene ID<sup>b</sup> | Gene ID<sup>b</sup> | Gene ID<sup>b</sup> | Gene ID<sup>b</sup> | Localisation<sup>d</sup> | Reference |
| HSP90-alpha/HSPC2<br>HSP90-beta/HSPC3 | 3324<br>3326 | Tb927.10.10890 |  | LmjF.33.0312 |  |  |  |  |
|  |  | Tb927.10.10900 |  | LmjF.33.0314 |  |  |  |  |
|  |  | Tb927.10.10910 |  | LmjF.33.0316 |  |  |  |  |
|  |  | Tb927.10.10920 | TcCLB.507713.30 | LmjF.33.0318 |  |  |  |  |
|  |  | Tb927.10.10930 | C4B63_113g25 | LmjF.33.0320 |  |  |  |  |
|  |  | Tb927.10.10940 | C4B63_113g29 | LmjF.33.0323 |  |  |  | (58) |
|  |  | Tb927.10.10950 | C4B63_113g30 | LmjF.33.0326 |  |  | CYT | (61) |
|  |  | Tb927.10.10960 | C4B63_113g33 | LmjF.33.0330 | CFAC1_280011900 | BSAL_87515 | NUC | (70) |
|  |  | Tb927.10.10970 | C4B63_84g87 | LmjF.33.0333 | CFAC1_280012000 |  | FLAGELLAR | (71) |
|  |  | Tb927.10.10980 | C4B63_84g88 | LmjF.33.0336 |  |  | CELL SURFACE | (87) |
|  |  | Tbg972.10.13260 | C4B63_84g89 | LmjF.33.0340 |  |  |  |  |
|  |  | Tbg972.10.13270 | Tc_MARK_3581 | LmjF.33.0343 |  |  |  |  |
|  |  | Tbg972.10.13280 |  | LmjF.33.0346 |  |  |  |  |
|  |  |  |  | LmjF.33.0350 |  |  |  |  |
|  |  |  |  | LmjF.33.0355 |  |  |  |  |
|  |  | Tb11.v5.0543* |  | LmjF.33.0360 |  |  |  |  |
|  |  |  |  | LmjF.33.0365 |  |  |  |  |
| **GRP94/HSPC4** | 7184 | Tb927.3.3580 | C4B63_10g439 | LmjF.29.0760 | CFAC1_100018800 | BSAL_88715 | ER | (58) |
|  |  | Tbg972.3.3850 | Tc_MARK_3058 |  |  |  | NUC | (61) |
|  |  |  |  |  |  |  | FLAGELLAR | (70) |
|  |  |  |  |  |  |  | CELL SURFACE | (71) |
| **TRAP-1/HSPC5** | 10131 | Tb927.11.2650 | TcCLB.504153.310 | LmjF33.2390 | CFAC1_230028300 | BSAL_33145 | MITO | (62) |
|  |  | Tbg972.11.2900 | C4B63_2g430 |  |  |  | FLAGELLAR | (70) |
|  |  |  | Tc_MARK_6238 |  |  |  |  | (87) |

<sup>a</sup> The nomenclature for the Hsp90/HSPC proteins from *T. b. brucei*, and *T. b. gambiense* were derived according to Folgueira and Requena (2007).
<sup>b</sup> The Gene IDs for the members of the *T. b. brucei* (Tb refers to Tbb), *T. b. gambiense*, *T. cruzi*, *C. fasciculata*, *B. saltans* and *L. major* Hsp90/HSPC protein family were retrieved from the TriTrypDB database (http://tritrypdb.org/tritrypdb/; (78). The Gene IDs for the members of the *H. sapiens* Hsp90/HSPC protein family were retrieved from NCBI (https://www.ncbi.nlm.nih.gov/).
<sup>c</sup> The Gene IDs for the orthologues, identified by reciprocal BLASTP analysis, of three strains of *T. cruzi* are listed. *T. cruzi* CL Brener Esmeraldo-like (TcCLB), *T. cruzi* Dm28c 2018 (C4B63), and *T. cruzi* marinkelli strain B7 (Tc\_MARK).
<sup>d</sup> Subcellular localizations for the *T. brucei* Hsp90/HSPC proteins were either acquired from using the TrypTag database (http://tryptag.org/; (87) and/or predicted using various proteomic datasets and online prediction software listed in the materials and methods.
CYT-Cytosol; MITO- Mitochondrion; NUC- Nucleus; ER- Endoplasmic reticulum; GYLCO- glycosomes; FLAGELLAR- Flagellar; CELL SURFACE- Cell surface.
\* Putative *T. b. brucei* HSP83 gene has not been assigned to a chromosome on the TriTrypDB database (http://tritrypdb.org/tritrypdb/; (78) yet.

Syntenic analysis was conducted to evaluate the conservation of the gene arrangement of the cytosolic Hsp83 genes in *T. brucei* and selected kinetoplastid parasites. The conserved syntenic regions surrounding the selected Hsp83 genes were searched by examining the conserved colocalization of neighbouring genes on a scaffold of the *T. brucei* subspecies (*T. b. brucei* and *T. b gambiense*) and selected kinetoplastid parasites for this study using genome information from TriTryDB. The identities of unknown neighbour genes of the selected Hsp83 genes were conducted using a BLASTP search on the NCBI database.

### Physiochemical properties, protein expression, and the determination of the organelle distribution for the *T. brucei* Hsp90/HSPC complement

The physiochemical properties, molecular weight (Da) and isoelectric point (pI) of each gene was determined using the compute pI/Mw tool from ExPASy (https://web.expasy.org/compute_pi/) (84). Data on the previously reported phenotypic RNAi knockdown screen, (85), for each member of the Hsp90/HSPC complement and identified Hsp83 co-chaperones were retrieved from TrypsNetDB (http://trypsnetdb.org/QueryPage.aspx) (86). The predicted organelle distribution for each protein was searched using the TrypTag microscopy project’s online server, (87). This project aims at tagging every trypanosome protein with mNeonGreen (mNG) (88) to determine the protein’s localization in the cell within the parasite (http://tryptag.org/)(87). Proteomic data from the mitochondrion (62), mitochondrial importome (63), respiratome (65), mitochondrial membranes (outer, intermembrane space, inner and matrix) (65), nucleus (60), nuclear pore (66), glycosomes (67, 68), flagellum (69, 70) and cell surface (71) were also used for the prediction of the organelle distribution for the *T. brucei* Hsp90 complements and Hsp90/ HSPC complements and Hsp83 co-chaperones.

### Identification of potential post-translational modification sites for the *T. brucei* Hsp83 proteins

Mass spectrometric information from a collection of relevant databases on *T. brucei* PTMs (73,75–77) for the relevant proteins was retrieved using the previously identified accession numbers. Information on the respective PTMs (modification sites, modification types and modified residue) were obtained and the modified residues were mapped onto Fig S1 for all Hsp90 isoforms from *T. brucei* subspecies (*T. b. brucei* and *T. b gambiense*) with orthologues from other kinetoplastids and from human, then analysed for determination of conserved and specific PTMs among the *T. brucei* Hsp90 complements.

## Results and discussion

### Determination of the *T. b. brucei* and *T. b. gambiense* Hsp90/HSPC complement

The protozoan parasite, *T. brucei* is comprised of three subspecies, with the genomes of *T. b. gambiense* and *T. b. brucei* already sequenced (89, 90). Any information obtained from the genome of the non-human infective *T. brucei* subspecies, *T. b. brucei*, can be inferred for the human infective subspecies, *T. b. rhodesiense*, as the *T. b. brucei* TREU927 strain displays the full range of known *T. brucei* phenotypes and possesses similar biological and genetic characteristics (90). However, the *T. b. gambiense* genome was sequenced due to the subspecies displaying profoundly different biological and genetic characteristics (89). An *in silico* analysis of the Hsp90/HSPC complement in both *T. brucei* subspecies was conducted to provide an overview of the *T. brucei* Hsp90 family. The nomenclature and format to categorize the *T. brucei* Hsp90 family was adopted from our previous study (9). The orthologue of the cytosolic Hsp90 member in *T. brucei* is termed Hsp83 (91), and thus will be referred to as Hsp83 in this study. This protein displays variable molecular weight amongst different kinetoplastid protists. However, to underscore whether discussing a protein from *T. b. gambiense* or *T. b. brucei*, the abbreviations Tbg and Tbb respectively, were used in this study. The orthologous relationships of the Hsp90 family from *T. b. brucei* and *T. b. gambiense* to the selected organisms in this study are presented in Table 1, and a comprehensive domain organisation of the predicted *T. brucei* Hsp90 proteins is illustrated in Fig 1.

**Fig 1.**
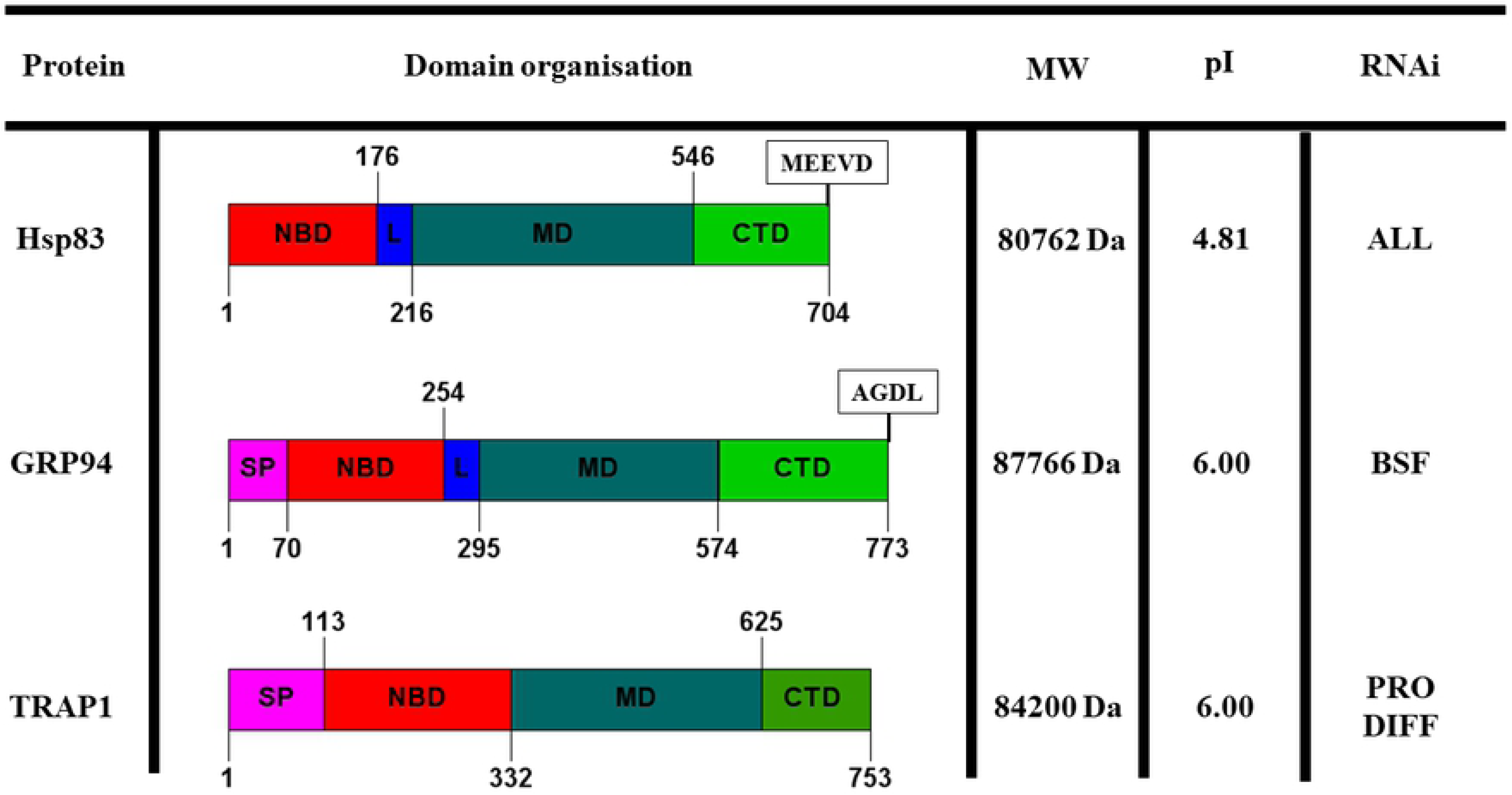
Schematic representation of the domain architecture of the Hsp90/HSPC proteins in *T. brucei*. Each protein sequence is represented by a coloured bar with the numbering on the bottom of the bar indicating the length of the protein in amino acid residues. Protein domains and other associated features that were identified using Prosite (80) and SMART (79) are also shown, and include the N-terminal nucleotide binding domain (NBD; red), variable charger linker domain (L; dark blue), middle client protein-binding domain (MD; light blue), a C-terminal dimerization domain (DD; green) and targeting signal peptides (SP; pink). The physiochemical properties, molecular weight (MW) and isoelectric point (pI), for each *T. brucei* Hsp90 protein was calculated using the compute pI/Mw tool from ExPASy (https://web.expasy.org/compute_pi/; (84). Data on the phenotypic knockdown screen using RNAi conducted by Alsford et al. (2011), for Hsp90/HSPC protein member is provided: ALL-required for all life cycle stages; BSF-required for bloodstream form; PRO-required for procyclic form; DIFF-required for differentiation; NE-Non-essential; ND-Not determined.

Twelve putative *Hsp90* genes were identified to be encoded on the *T. b. brucei* genome (Table 1), which is consistent with previous findings (49, 91), while *T. b. gambiense* was identified in this study to only have 5 putative *Hsp90* genes encoded on its genome. The reduction in the *Hsp90* gene numbers found in *T. b. gambiense* could be a consequence of the reduced genome size observed in the human infective subspecies (92). The intraspecific genomic variation is largely associated with tandem or segmental duplications observed in *T. b. brucei* (89). This study also identified a new unassigned putative *Hsp90* gene (Tb11.v5.0543) in the animal infective subspecies, *T. b. brucei*. Though, whether this gene represents an additional and/or novel *Hsp90* gene needs to be further verified (Table 1). For the putative Hsp90 genes identified in this study for *T. b. brucei*, 10 of the 12 putative *Hsp90* genes identified were found to be homologous to Hsp83, whereas in *T. b. gambiense*, 3 of the 5 putative *Hsp90* genes identified were homologous to Hsp83 (Table 1). The remaining two *Hsp90* genes found in both *T. b. brucei* (Tb927.3.3580 and Tbg972.3.3850) and *T. b. gambiense* (Tb927.11.2650 and Tbg972.11.2900) showed significant identity to the ER and mitochondrial resident paralogues of Hsp90, GRP94 and TRAP-1 respectively (Table 1). This indicates that a single gene copy for GRP94 and TRAP-1 is encoded on the genome in both *T. brucei* subspecies. Phylogenetic analysis shows that the *T. brucei* Hsp90/HSPC family is also comprised of 3 distinct Hsp90 groups (Hsp83, GRP94 and TRAP-1), which cluster into clades according to protein sequence and subcellular localisation (Fig 2). In contrast to humans with 4 Hsp90 isoforms, there are 3 Hsp90 isoforms (Hsp83, GRP94 and TRAP-1) identified by phylogenetic analysis to be present in all kinetoplastid organisms used in this study (Table 1; Fig 2).

**Fig 2.**
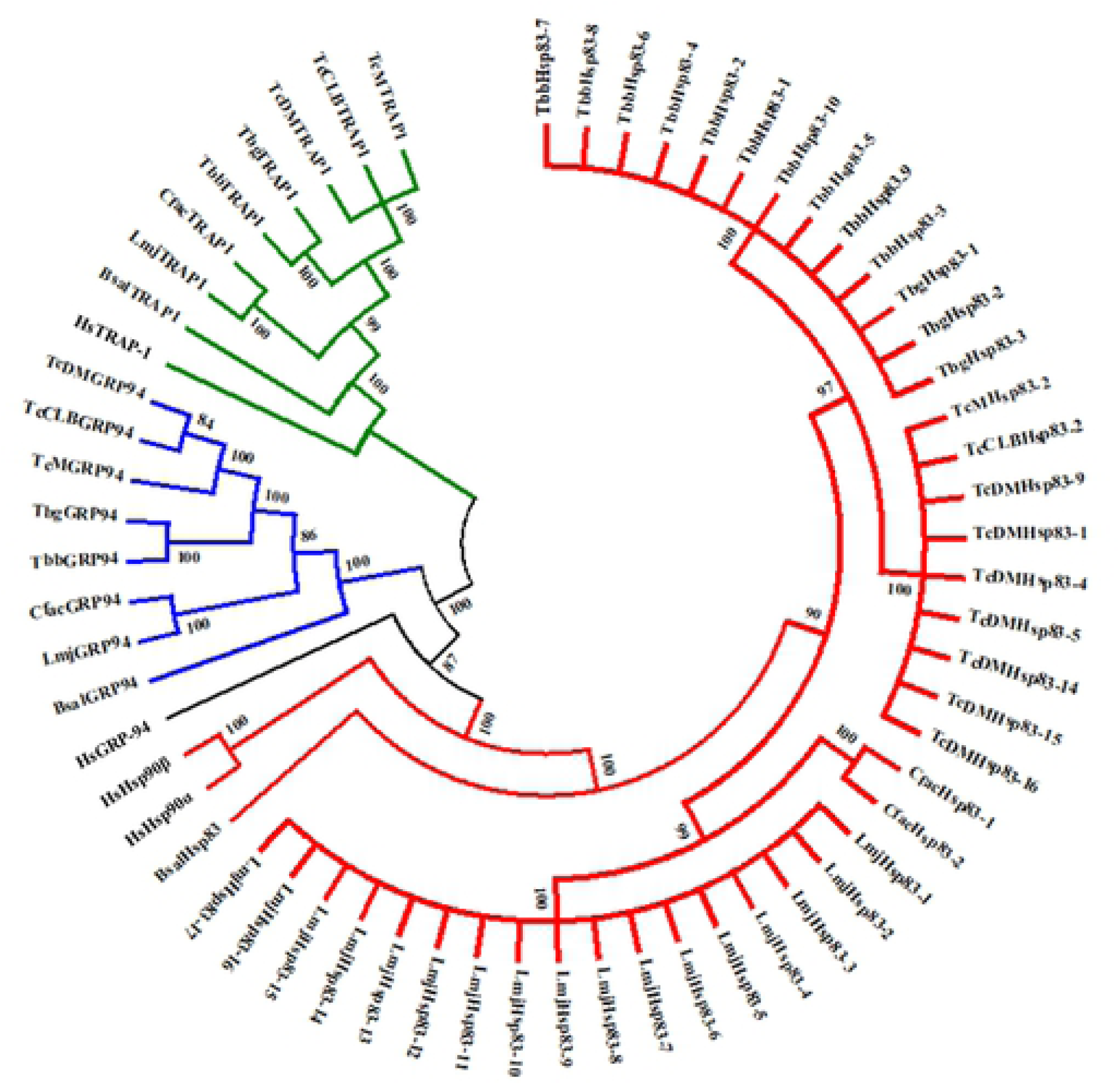
Phylogenetic analysis of the Hsp90/HSPC family from *T. brucei* in relation to human and selected kinetoplastid parasites. Multiple sequence alignment of the full-length amino acid sequences of the Hsp90/HSPC gene families in human and selected kinetoplastid parasites. The multiple sequence alignment provided in Fig. S1 was performed using the in-built ClustalW program (81) with default parameters on the MEGA X software (82). The phylogenetic tree was constructed by MEGA X using the Maximum-likelihood method based on the Jones–Taylor–Thornton (JTT) matrix-based model of amino acid substitution (83) with gamma distribution shape parameter (G). The alignment gaps were excluded from the analysis, and the number of amino acid sites used to construct the tree numbered 572. Bootstrap analysis was computed with 1000 replicates. Gene ID/Accession numbers for the *T. b. brucei* (Tbb), *T. b. gambiense* (Tbg), *T. cruzi* (TcCLB, CL Brener Esmeraldo; TcM, marinkellei strain B7; TcD, Dm28c 2018), *C. fasciculata* (Cf), *B. saltans* (Bs), *L. major* (Lmj) and human (Hs; *H. sapiens).* Hsp90 amino acid sequences can be found in Table S1. The subcellular localisation for Hsp90s is indicated by coloured branches. Red: cytosolic; blue: endoplasmic reticulum; and green: mitochondrion. Scale bar represents 0.2 amino acid substitutions per site.

Previous literature reported that 11 *Hsp90* genes are encoded on the *Trypanosoma cruzi* (*T. cruzi*) genome (50). In this study we included three different *T. cruzi* strains: CL Brener Esmeraldo-like (TcCLB), Dm28c 2018 (C4B63), and marinkelli strain B7 (Tc_MARK) to determine the Hsp90/HSPC in the American trypanosome. It was identified in this study that the *T. cruzi* CL Brener Esmeraldo-like strain encodes 2 *Hsp90* genes, the Dm28c 2018 strain has 9 *Hsp90* genes, and the marinkelli strain B7 has 3 *Hsp90* genes (Table 1). However, it was found in this study that many of the *Hsp90* genes homologous to Hsp83 identified in the three *T. cruzi* strains were partial and/or truncated genes. These genes may be the products of non-sense mutation leading to premature termination of Hsp83, which if expressed in the parasite can code for truncated Hsp83 proteins. These partial and/or truncated *Hsp83* genes in this study were omitted from the analysis. The obvious discrepancy in numbers of genes amongst the *T. cruzi* strains, and its numerous partial and/or truncated Hsp90 sequences calls for re-evaluation of its genome annotation. *Leishmania major* (Lmj) contains the largest Hsp90 family with a total of 19 *Hsp90* genes, 17 of which were found to be homologous to Hsp83, and these findings are like previous studie**s** (8,49,50). Other kinetoplastids included in this study were the non-parasitic *Bodo saltans* (*B. saltans*) (93) and the insect infecting *Crithidia fasciculata* (*C. fasciculata*) (94), which were found to encode 3, and 4 putative *Hsp90* genes respectively (Table 1). Both these kinetoplastids were found to possess genes encoding for all three Hsp90 isoforms (Hsp83, TRAP-1 and GRP94), though *C. fasciculata* was identified to possess two *Hsp83* gene (Table 1). Early genomic studies suggested that the human genome contained 16 Hsp90/HSPC genes (5 functional and 11 pseudogenes), which have been categorised, according to the proposed standardized guidelines for HSP nomenclature, into 4 isoforms under the superfamily name HSPC (12, 21). In contrast to the kinetoplastid protists, humans have two isoforms of Hsp90 localized in the cytoplasm: the inducible form Hsp90α/HSPC2 and the constitutive form HSP90β/HSPC3 (20). *T. brucei* Hsp83 is 62% and 63% identical at the amino acid level to HsHSPC2 and HsHSPC3 respectively, and this sequence identity increases to over 70% in the NBD (Fig S1). Phylogenetic analysis has suggested that the two cytosolic isoforms (heat-induced Hsp90α/HSPC2/HSP90AA2 and constitutively expressed Hsp90β/HSPC3/HSP90AB1) arose from gene duplication, and the organelle Hsp90s (GRP94/HSPC4 and TRAP-1/HSPC5) developed from a common ancestor (95–97). Hsp83 (Tb927.10.10980) and TRAP-1 (Tb927.11.2650) were identified as phosphoproteins in this study, while kinases are yet to be identified in the ER and little is known about the effect of post-translational modifications on GRP94 (23, 98).

### Hsp83

Hsp83 and has been found to be an essential and highly abundant protein, that is encoded by multiple gene copies organized in a head-to-tail tandem array (49). It has been identified in this study and previous studies (49, 91) that *T. b. brucei* has been shown to encode for 10 tandem copies of *Hsp83* (Fig 3), whereas *T. b. gambiense* genome encodes 3 tandem copies of *Hsp83* (Fig 3). Syntenic analysis revealed that the *TbbHsp83* and *TbgHsp83* genes are both located on chromosome 10 in a head to tail orientation, with the same genomic organisation being observed in both *T. brucei* subspecies (Fig 3.). Like *T. brucei*, a discrepancy in *Hsp83* gene copy numbers was also observed for the three *T. cruzi* strains used in this study (Fig 3). Syntenic analysis revealed that the *T. cruzi* Dm28c 2018 (C4B63) strain has 16 tandem copies of Hsp83, though 9 were partial sequences (Fig 3), whereas both the CL Brener Esmeraldo-like (TcCLB) and marinkelli strain B7 (Tc_MARK) encode for 2 *Hsp83* genes, with 1 partial gene each (Fig 3). The genomes of the three *T. cruzi* strains need to be further investigated to determine if the partial sequences of the *Hsp83* genes are due to sequencing errors or a result of non-sense mutation. *L. major* has the highest *Hsp83* gene copy number with 17 tandem copies (Table 1; Fig 3), correlating with the high abundance of the protein being observed in *L. major* and several other *Leishmania* spp. (99). Syntenic regions surrounding the *Hsp83* genes were found to be virtually conserved across the selected kinetoplastids, with *B. saltans* being the exception (Fig 3). Thus, the discrepancy in gene copy number of Hsp83 in the two *T. brucei* subspecies and amongst the kinetoplastid organisms may have arisen from the differences in the life cycle of the kinetoplastids. Datamining of proteomic data revealed that all identified TbbHsp83 (TbbHsp83-1) proteins are present in both life cycle stages of the parasite: the bloodstream stage (BSF) and procyclic stage (PF) (58, 61). Though, the protein expression of the TbbHsp83 proteins were reported to be up regulated during at the BSF stage (58), despite gene regulation being unchanged in both the bloodstream and procyclic life cycle stages (61). All TbbHsp83 proteins were also present in the cell surface proteome (70) and TbbHsp83-10 (Tb927.10.10980) was found in the flagellar proteome (71).

**Fig 3.**
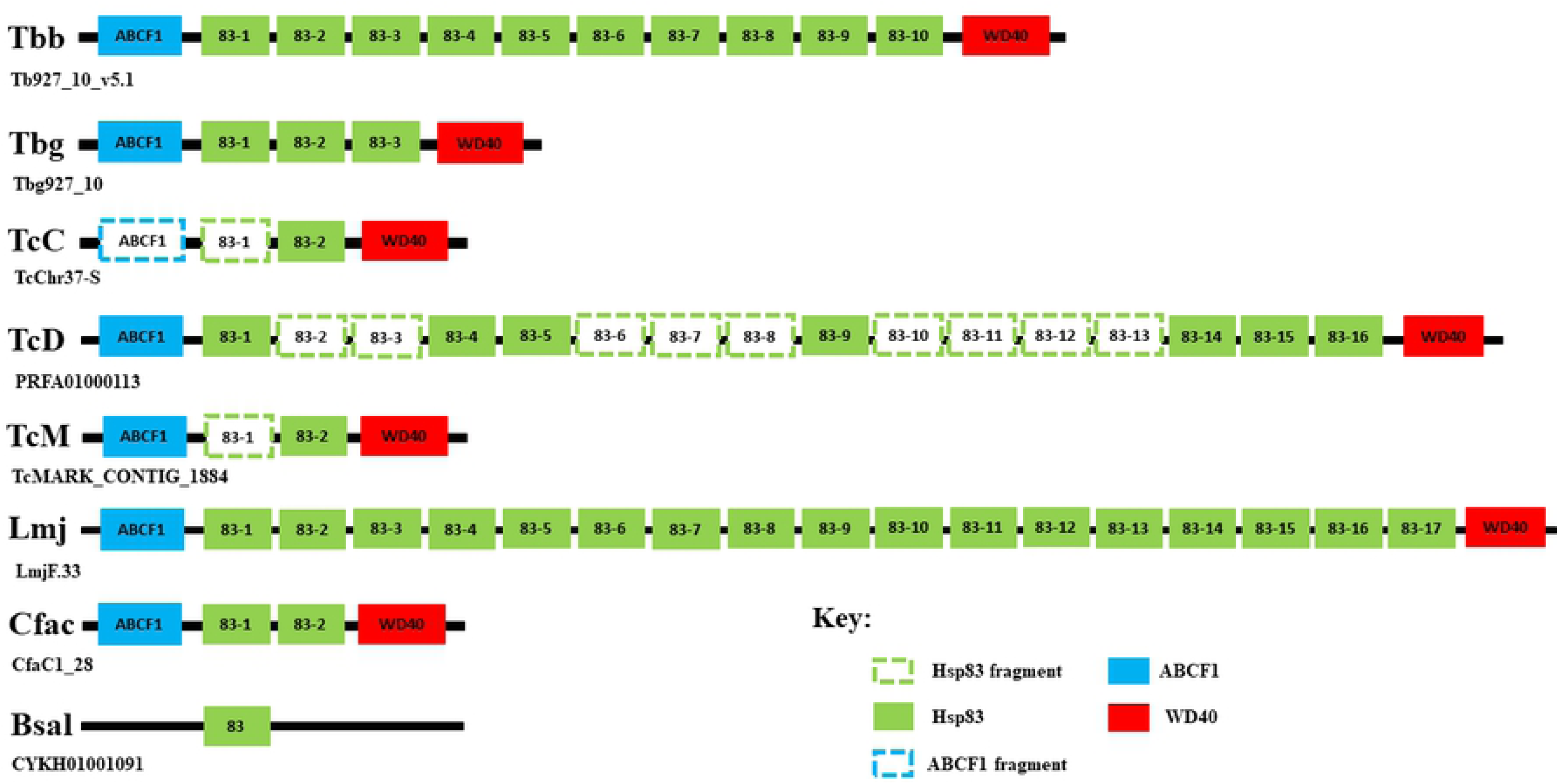
Syntenic analysis of the gene arrangement of the Hsp83 genes in *T. brucei* and selected kinetoplastid parasites. The conserved syntenic regions surrounding the selected Hsp83 genes were searched by examining the conserved co-localization of neighbouring genes on a scaffold of the *T. brucei* subspecies (*T. b. brucei* and *T. b. gambiense*) and selected kinetoplastid parasites : *T. cruzi* CL Brener Esmeraldo-like (TcCLB), *T. cruzi* Dm28c 2018 (TcD) strain, *T. cruzi* marinkelli strain B7 (TcM), *L. major* (Lmj), *B. saltans* (Bsal) and *C. fasciculata* (Cfac). The genome information used for this study was acquired from TriTrypDB database (http://tritrypdb.org/tritrypdb/; (78). The identities of unknown neighbour genes of the selected Hsp83 genes were conducted using a BLASTP search on the NCBI database. Abbreviations: ABCF1: ATP-binding cassette sub-family F member 1; WD40: WD40-repeat protein.

The amplification of HSP genes in protozoan parasites has been reported previously (8,9,51,100), and is considered a means by which the parasites increase chaperone levels in order to maintain proteostasis under normal and stressful conditions (53). The heat shock response is a highly conserved transcriptional program that in most organisms involves increased heat-shock gene transcription (101). However, in kinetoplastids, control of gene expression occurs almost exclusively at the post-transcriptional level, and that HSP synthesis during heat shock depends on regulation of mRNA turnover and translational control (102, 103). In *T. brucei*, post-transcriptional regulation of chaperone mRNAs is facilitated by a zinc finger protein, ZC3H11(104). The mRNA transcript levels of TbbHsp83 in BSF parasites increases >2-fold after heat shock (105), and is stabilized by ZC3H11 to promote the survival of the parasite (104). Treatment of *T. b. brucei* BSF parasites with 17-AAG sensitized the parasites to heat shock, as well as caused severe morphological abnormalities and cell cycle disruption (46). Pharmacological inhibition of Hsp83 activity in several *Leishmania* spp. Induced morphological and biochemical promastigote-to-amastigote differentiation (53,106,107), which mimics environmental triggers such as heat shock and acidic milieu, indicating a pivotal role for Hsp83 in kinetoplastid protists in environmental sensing and life cycle control. Interestingly, treatment of *T. cruzi* bloodstream trypomastigotes with geldanamycin induced morphological changes in the parasites but not life cycle progression (44). Therefore, Hsp90 cellular homeostasis as a key factor for the control of stage differentiation appears to be dependent on the tropism of the parasite and the different regulatory pathways for life cycle control. It would be interesting to investigate if the pharmacological inhibition of Hsp83 effects cellular differentiation amongst the three *T. brucei* subspecies.

The monophyletic cluster of the cytosolic Hsp83s suggests a general conservation of function, structure, and sequence in the kinetoplastid Hsp83 homologues (Fig 2). The 704 amino acid sequences of the corresponding TbbHsp83 and TbgHsp83 proteins were found to be almost identical (Fig S1) and contain the predicted canonical domain architecture of typical cytosolic Hsp90s (Fig 1). Despite the overall structural conservation, kinetoplastid Hsp83 proteins possess unique biochemical features which separate them from their human counterparts and can be potentially exploited in selective drug discovery studies. Unlike mammalian Hsp90, which binds ATP but has low basal ATPase activity, TbbHsp83 from *T. b. brucei* shows potent ATPase activity (108). This enhanced ATPase activity has also been shown in Hsp83 orthologue in *T. cruzi* (109). As a consequence of its greater affinity for ATP, TbbHsp83 also possesses a greater affinity for Hsp90 NBD specific inhibitors, and thus is more sensitive to inhibition by these compounds than its human orthologues (46, 108). In a study conducted by Pizarro and colleagues (2013), biophysical and biochemical techniques were able to identify three short divergent regions in the TbbHsp83 NBD, that they targeted for selective pharmacological inhibition of TbbHsp83 over human Hsp90. The higher ATPase activity is predicted to be a result of the parasites enhanced requirement for proteostasis maintenance by molecular chaperone through stabilizing key cell regulators under hostile conditions (108). This observation is consistent with human Hsp90’s pivotal role in some forms of cancer through conformational regulation of labile kinases and ligases (19,41,43,110).

It was also interesting to note that the variable charged linker domain which links the NBD to the MD in cytosolic Hsp90s found in higher eukaryotes is also present in *T. brucei* (Fig 1). This region is highly divergent in both length and amino acid sequence among Hsp90 proteins of different species and does affect Hsp90 function, co-chaperone interaction, and conformation (111–113). Hsp90 from protozoa often have extended linkers with the malaria parasite, *Plasmodium falciparum*, exhibiting one of the longest linkers reported thus far (45). Complementation of human and yeast charged linkers by the *P. falciparum* version reduces ATPase activity and affects client protein binding (113), thus indicating that this linker could provide specificity to the activity of Hsp90 from different species. Therefore, comparative analysis of *T. brucei* Hsp83 proteins with their human counterparts, as well as linker swapping experiments, will be especially useful in understanding the role of the linker region in *T. brucei* Hsp90 biology, and possible future exploitation as a unique drug binding region.

Post-translational modifications, and particularly phosphorylation of tyrosine, serine, and threonine residues at multiple sites of cytosolic Hsp90 is a well-known chaperone activity modulator mechanism in many organisms (114–117). Hsp90 steady-state phosphorylation is species-specific relative to the different cellular environments (116). S53 and S286 were determined to be phospho-modified residues that were conserved within the ten cytosolic *T. brucei* Hsp83 proteins, while T211, T216, S597 and S694 were conserved in all analysed kinetoplastids in this study (Fig. S1). S374 was conserved in both kinetoplastids and humans (Fig. S1). The same phospho-modified residues were previously described for the cytosolic Hsp83 orthologue from *L. donovani* (117). The following acetylation sites were predicted for TbHsp83: K44, K227, K277, K259, K289, K337, K394, K418, K421, K474, K487, K515 and K533. The residues conserved amongst the other isoforms were mapped in Fig. S1. The predicted N-glycosylation sites, N90, N372 and N612 were conserved in all kinetoplastids and humans, whilst N51 was determined to be specific to *T. brucei* Hsp83 (Fig. S1). Two ubiquitination sites (K394 and K560) were found conserved in all analysed cytosolic Hsp90 isoforms in this study (Fig. S1).

### TRAP-1

The mitochondrial isoform of the Hsp90/HSPC family was first identified in association with the mammalian tumour necrosis factor 1 (TNF-1) protein, hence termed TRAP-1 (118). It was promptly suggested as a member of the 90-kDa molecular chaperone family due to strong homology with other Hsp90 members (118). Since then, TRAP-1/HSPC5 orthologues have been identified in a variety of eukaryotic and prokaryotic organisms. This study identified a single entry for a putative *TRAP-1* gene annotated in the genomes of both *T. b. brucei* (Tb927.11.2650) and *T. b. gambiense* (Tbg972.11.2900) (Table 1). The selected kinetoplastids in this study also encoded a single copy of TRAP-1 (Table 1) which was consistent with previous studies (49), except for *T. cruzi* which was previously stated to encode for two TRAP-1 orthologues (49, 50). Phylogenetic analysis indicates a general conservation of kinetoplastid TRAP-1 (Fig 2), though little experimental characterization of these genes has been conducted in kinetoplastids. It is predicted that the cellular role of the kinetoplastid TRAP-1 proteins will be orthologous to HsHSPC5, whose major function is to maintain mitochondrial integrity, modulate mitochondrial metabolism and protect against mitochondrial apoptosis (24). Furthermore, HSPC5 counteracts protein aggregation inside the mitochondria and supports protein folding (119), leading to healthy, intact mitochondria.

Mammalian TRAP-1 orthologues are localized predominantly in the mitochondrial matrix, where at least 6 different protein variants were found resulting from differing splicing patterns, amino acid additions and/or deletions (120, 121). The translation of the main TRAP-1 mRNA generates a precursor protein of 704 amino acids, that contains a putative 59-amino acid, N-terminal mitochondrial import sequence which is removed upon organelle import (121, 122). It was predicted that both TbbTRAP-1 and TbgTRAP-1 localize in the mitochondria, as the proteins possess a positively charged N-terminal leader sequence (Fig 1). Proteomic and localisation studies confirmed that TbbTRAP-1 localises to the mitochondria (62, 87), the protein was also detected in the flagella of *T. b. brucei* BSF parasites (70) (Table 1). The subcellular distribution of TbbTRAP1 during the parasite’s life cycle could be related to the shape and functional plasticity of the *T. brucei* single mitochondrion, which undergoes profound alterations to adapt to the different host environments (123). Phenotypic knockdown of TbbTRAP-1 had a detrimental effect on the survival and fitness of the parasite at the procyclic stage of its life cycle and negatively affected parasite differentiation (85). Thus, *T. brucei* TRAP-1 proteins may be an important modulator of mitochondrial bioenergetics at the procyclic stage, as well play an integral role in parasite pathogenesis.

In terms of PTMs, 3 putative phosphorylation sites were found in the middle domain of TRAP-1. S286 and S363 were specific phosphorylation sites for TRAP-1 and S374 was conserved amongst all the Hsp90 proteins (Fig. S1). Several amino acids have been reported as potential targets for post-translational modifications in human TRAP-1, yet its phosphorylation mechanism remains to be revealed (24). K109, K480 and K601 were predicted to be specific acetylation sites for TbTRAP-1. Additional TbTRAP-1 putative acetylation sites on lysine residues were conserved amongst the mitochondrial isoforms from all analysed taxon (Fig. S1). Most of these PTMs of Hsp90 and other inferences stated here are yet to be verified experimentally

### GRP94

The glucose-regulated 94 kDa protein (GRP94) is a Hsp90 family member residing in the lumen of the endoplasmic reticulum (ER) (98), where it is involved in the maturation of membrane-resident and secreted protein clients (23). GRP94 is present as a single gene in all metazoa, although the gene is not found in many unicellular organisms such as bacteria, archaea, yeast, and most fungi (23). This study identified a single putative entry for the *GRP94* gene in both *T. brucei* subspecies and the selected kinetoplastid protists (Table 1). These findings are consistent with previous findings for *T. brucei* and *L. major* (49), though previous reports indicated that *T. cruzi* CL Brener Esmeraldo-like strain encodes 3 GRP94 orthologs (49, 50). The genome of the *T. cruzi* strain needs to be further investigated to determine if these partial sequences of the *GRP94* genes (TcCLB.506591.4 and TcCLB.503811.10) are due to sequencing errors.

Both TbbGRP94 and TbgGRP94 genes are present on chromosome III and encode proteins considerably longer in amino acid sequence when compared to Hsp83 (Fig 1), which is characteristic of GRP94 protein members (13, 124). GRP94 proteins share structural similarity with cytosolic Hsp90 proteins, though the N-terminus contains an ER signal peptide while the C-terminal MEEVD peptide is replaced with the KDEL motif that is required for retention in the ER (98). Sequence analysis of TbbGRP94 and TbgGRP94 indicates that the GRP94 proteins share domain architecture with typical GRP94 proteins including the possession of an N-terminal ER signal peptide (Fig 1). However, a variation in the C-terminal ER retention motif, KDEL, is observed in all the kinetoplastid orthologues of GRP94; AGDL in *Trypanosoma* spp., KEEL in *B. saltans*, EGDL in *C. fasciculata* and all *Leishmania* spp (Fig S1). Phylogenetic analysis indicates that the GRP94 proteins in kinetoplastid protists could have evolved separately from their mammalian orthologues (Fig 2), perhaps to fulfil a specific role within the parasites. Proteomic studies confirm the presence of GRP94 in flagella and cell surface (70, 71).

In kinetoplastids, the first recognized and characterized *GRP94* gene was in *Leishmania infantum* (*L. infantum*). The GRP94 orthologue in *Leishmania infantum* (*L. infantum*) was shown to localise in the ER and shares many of the activities of GRP94s of other eukaryotes (125). Unlike GRP94 in mammalian cells, LinGRP94 is not essential for cell viability and *LinGRP94* mRNA is induced developmentally rather than by canonical GRP94-inducing stresses (125). The protein was highly immunogenic during *Leishmania* infection (126, 127), and essential for lipophosphoglycan (LPG) assembly (125), an abundant surface glycolipid of *Leishmania* promastigotes that is critical to parasite virulence (128). Effectively, the critical role of GRP94 in *Leishmania* appears to be adapted to the synthesis of glycoconjugates and directing the host immune response implicating a pivotal role in parasite virulence (125). However, whether this specialized role is conserved in *T. brucei* and other kinetoplastid parasites will need to be elucidated. The function and cellular roles of TbGRP94 should be explored, given the immunogenic and antigenic properties shown by the *L. infantum* GRP94, as this protein could constitute a valuable molecule for diagnostic purposes, and quite possibly a potential candidate for studies of protective immunogenicity.

### The *T. brucei* Hsp83 co-chaperone system

In all organisms, Hsp90 is a dynamic protein that undergoes a conformational cycle whose directionality is determined in large part by ATP binding and hydrolysis, together with a cohort of co-chaperones (35,129,130). The Hsp90 chaperone ensembl can vary in composition depending on the client proteins, but usually includes Hsp70/J-protein, p23, immunophilins, Aha1 and STIP1 (HOP) (130). The variation in subunit composition across organisms appears to be related to the fact that the function of some Hsp90 co-chaperones may be restricted to specific subsets of client proteins, be required for client protein activation in a species-dependent manner, or made redundant by other co-chaperones (36). The Hsp90 chaperone system in intracellular protozoan parasites has been explored in previous studies (55, 131). Thus, using the human and kinetoplastid systems, this study analysed the composition of the *T. brucei* Hsp83 chaperone system. It was determined in this study that *T. brucei* possesses an almost complete set of co-chaperones (Table 2), with the only notable absence being cell division cycle 37 (Cdc37). The absence of a gene encoding for Cdc37 has also been noted in several intracellular protozoan parasites (55,117,132,133) and was not evident in 10/19 species examined by a study conducted by Johnson and Brown (2009). Cdc37 is a co-chaperone that has a specialized and indispensable role in the maturation and/or stabilization of a large subset of protein kinases (134). The absence of Cdc37 in some species shows that clients that are dependent on a specific cochaperone in one species may not require Hsp90 for function in other species, thus the protein kinases in protozoan parasites may have evolved in such a way that the proteins bind a different co-chaperone or are independent of Hsp90 for function. Since little is known about why a protein becomes dependent on Hsp90 for activity or stability, it poses interesting questions on the mechanism by which the maturation and regulation of protein kinases in protozoan parasite is mediated dependent or independent of Hsp83. Exploration of this mechanism may provide a potential avenue for chemotherapeutics since protein kinases are also an attractive drug target in infectious disease, such as African Trypanosomiasis. The Hsp70/J-protein machinery from *T. brucei* have been explored previously (9). The identified Hsp83 co-chaperones in both *T. brucei* subspecies are listed in Table 2, and a comprehensive domain organisation of these predicted proteins is illustrated in Fig 4. Additionally, the Hsp83 co-chaperones were categorised in this study based on the presence of the TPR domain.

**Fig 4.**
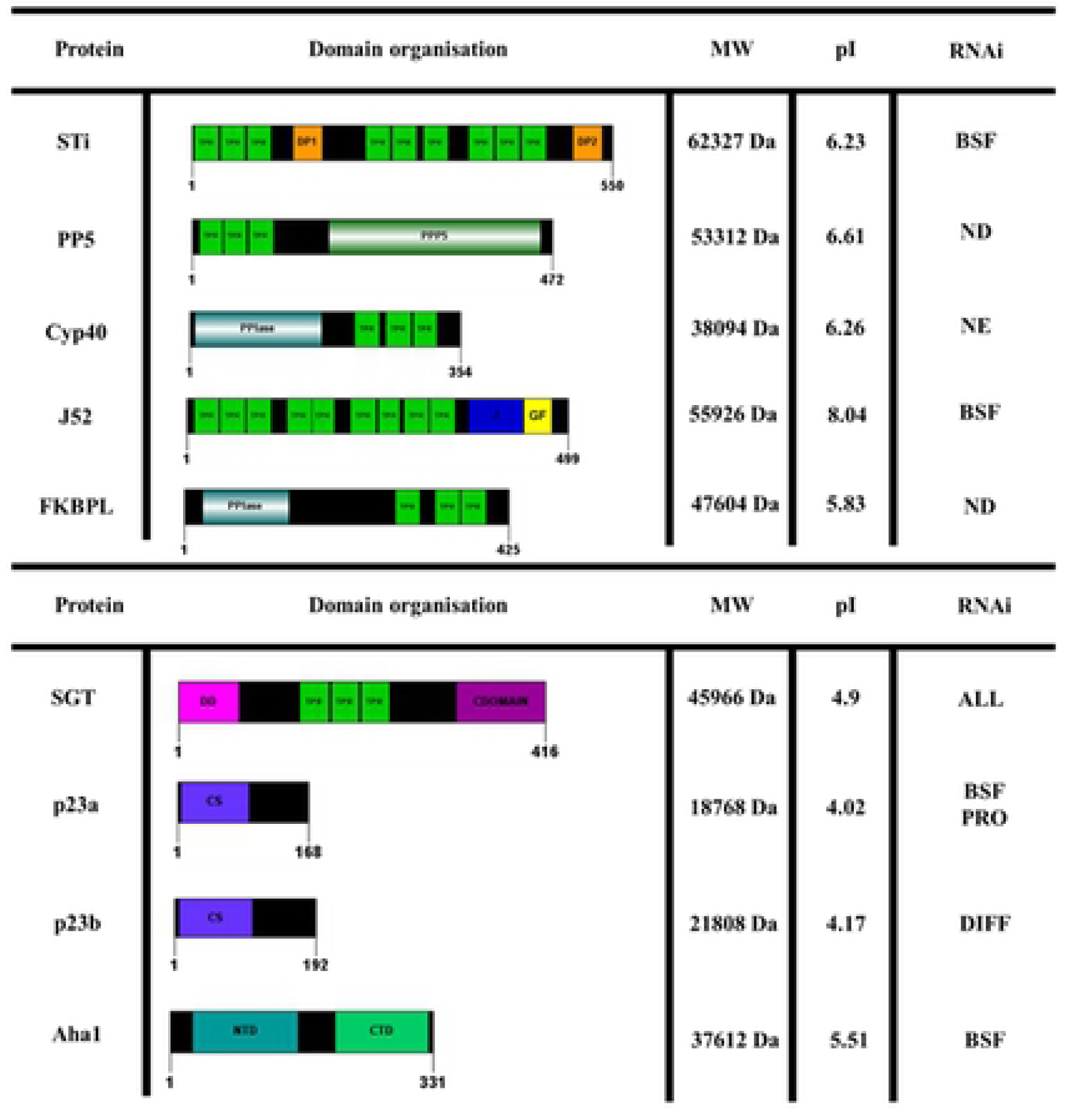
Schematic representation of the domain architecture of the Hsp83 TPR and non-TPR co-chaperones in *T. brucei*. Each protein sequence is represented by an open bar with the numbering on the bottom of the bar indicating the length of the protein in amino acid residues. Protein domains and other associated features that were predicted using Prosite (80) and SMART (79) are also shown. The physiochemical properties, molecular weight (MW) and isoelectric point (pI), for each *T. brucei* Hsp83 co-chaperone was calculated using the compute pI/Mw tool from ExPASy (https://web.expasy.org/compute_pi/; (84). Data on the phenotypic knockdown screen, using RNAi conducted by Alsford et al. (2011), for the Hsp83 co-chaperones are provided: ALL-required for all life cycle stages; BSF-required for bloodstream form; NE-Non-essential; ND-Not determined.

**Table 2.**
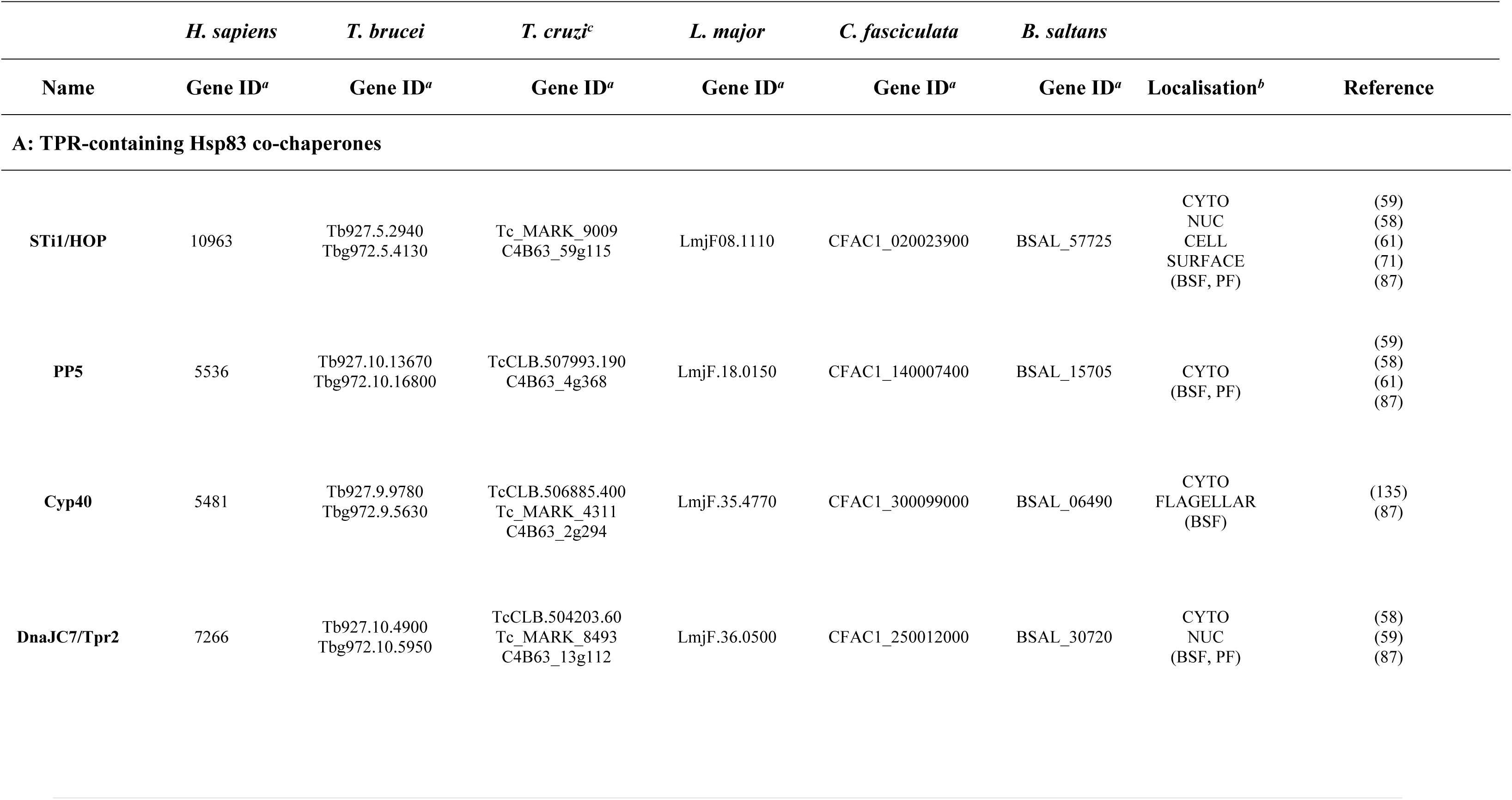
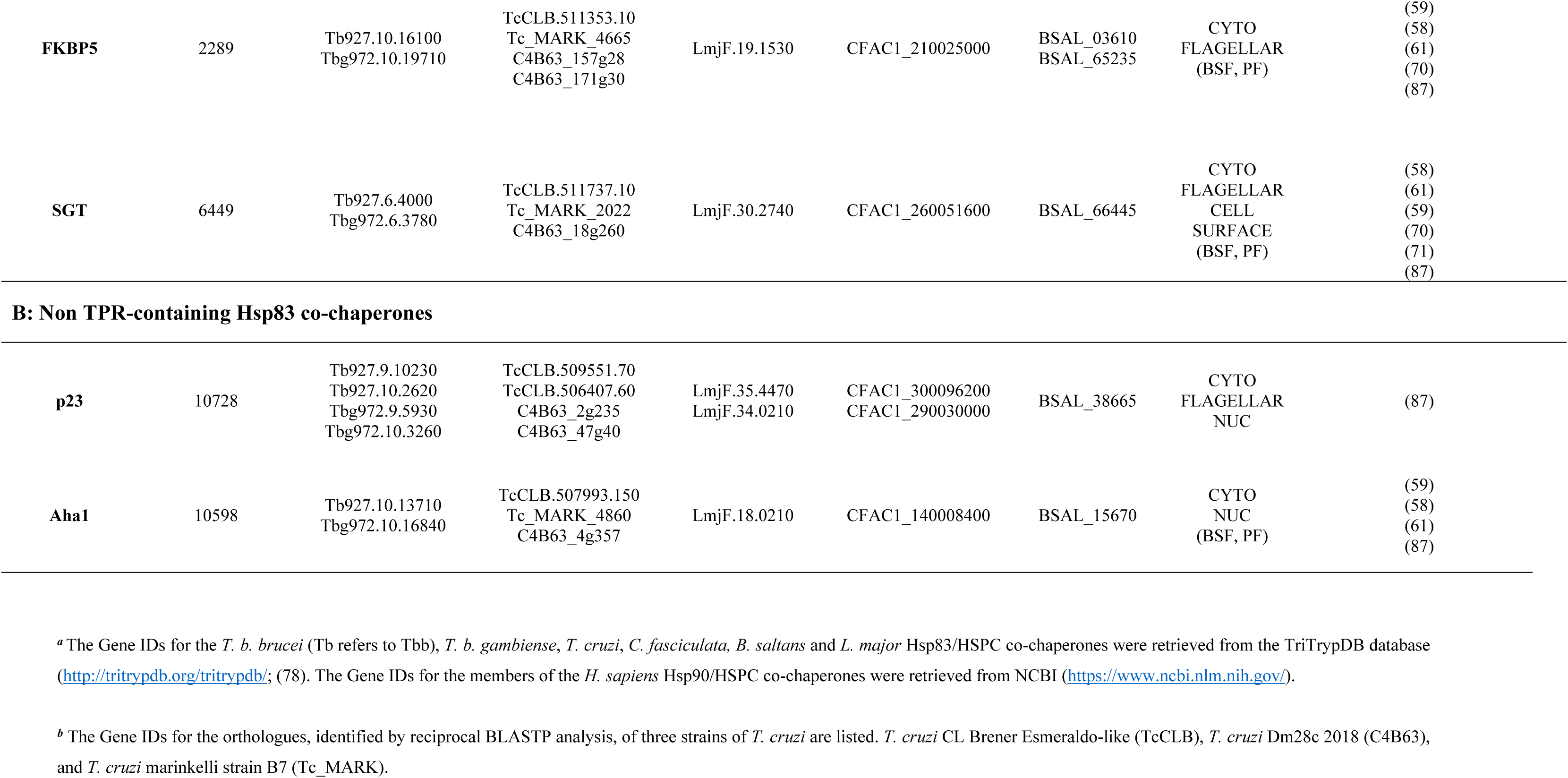
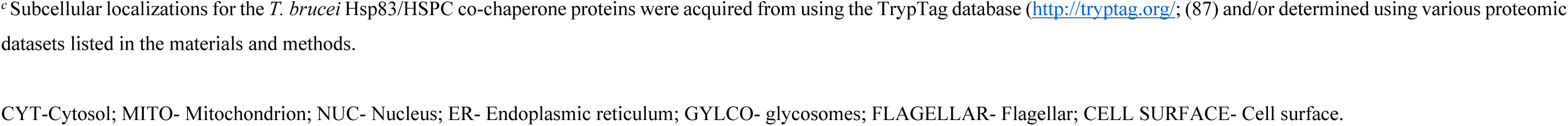
The Hsp83/HSPC co-chaperones from *Trypanosoma brucei* with their putative orthologues in *T. cruzi*, *L. major*, *C. fasciculata*, *B. saltans* and *H. sapiens*.

### TPR-containing co-chaperones

#### Sti1

Stress-inducible protein 1 (STI1), also known as Hsp70/Hsp90 organizing protein (HOP or STIP1) in mammals, is one of the best studied co-chaperones in the Hsp90 reaction cycle (136, 137) as it acts as an adaptor protein, mediating the interaction between Hsp70 and Hsp90 through its TPR domains (138–140). STI1/HOP is a widely conserved Hsp90 co-chaperone and has been annotated and characterized across diverse organisms including several kinetoplastid protists. Initially thought to be an indispensable protein, recent discoveries in yeast and some eukaryotes show that direct interaction can take place in vitro between Hsp70 and Hsp90 in the absence of HOP (141, 142). A single *STI/HOP* gene was found encoded in both *T. brucei* subspecies (Table 2), with the amino acid sequence indicating canonical STI/HOP domain architecture (Fig 4). Nine TPR motifs arranged into three TPR domains (TPR1, TPR2A and TPR2B) in addition to two domains rich in proline and aspartic acid (DP1 and DP2) were predicted (143, 144). Both STi/HOP orthologues in *T. cruzi* and *L. major* were found to immunoprecipitate with Hsp83 and Hsp70 as well as co-localize with these chaperones in the cytoplasm and/or around nucleus (145, 146). The expression of HOP isoforms was increased in response to different environmental stresses (145, 146) with LmjHOP being up regulated when the parasites are exposed to heat stress conditions (145), whereas only nutritional stress induced expression of TcSTi in the late growth phase of epimastigotes (146). The Hsp90-STi1 complex in *L. major* and *T. cruzi* has been shown to be pivotal to parasite differentiation (107, 145). Proteomic analysis in *Trypanosoma brucei* indicates that TbbSti1 is part of the cell surface (PF) proteome during the procyclic stage (71). Though TbbSti1 is present in both BSF and PF stages of the parasite, it was more highly expressed in the bloodstream form (58,59,61). These data suggest that the STi1 orthologue in both *T. brucei* subspecies should function as an adaptor protein for TbHsp83 and TbHsp70s, participating in the foldosome apparatus necessary for maintaining proteostasis, cytoprotection and modulating parasite differentiation.

#### PP5

Protein phosphatase 5 (PP5) is a member of the PPP family of serine/threonine protein phosphatases and it associates with Hsp90 in complexes during client protein maturation (147–149). PP5 is characteristically unique from other PPP family members in that it possesses an N-terminal TPR domain (150), which mediates interaction with Hsp90 (151). This interaction enables PP5 to modify the phosphorylation status of Hsp90 client proteins (149). The gene for PP5 in *T. b. brucei* (TbbPP5) has been extensively studied. TbbPP5 encodes a ∼52-kDa protein that possesses the canonical N-terminal TPR domain and phosphatase catalytic domain (152) as shown in Fig 4. TbbPP5 interacted with TbbHsp83 *in vivo* and co-localized with the chaperone in the cytosol of PRO parasites (56). Both TbbPP5 and TbbHsp83, upon heat shock and geldanamycin treatment, accumulated in the nucleus (56), indicating that both TbbPP5 and TbbHsp83 translocate to the nucleus when the parasites are exposed to proteotoxic stresses (56). TbbPP5 was detected in both BSF and PF stages of the parasite but upregulated in the procyclic form (58,59,61). Overexpression of TbbPP5 was found to partially negate the effect of geldanamycin treatment on cell growth, which indicates that the co-chaperone enhances the chaperoning function of TbbHsp83 and promotes the folding and maturation process of important regulatory molecules, which facilitate cell growth.

### Peptidyl-prolyl cis-trans isomerases (PPIases)

The immunophilin superfamily consists of highly conserved proteins with rotamase or peptidylprolyl cis-trans-isomerase (PPIase) activity that accelerates protein folding by mediating the isomerization of X-Pro-peptide bonds (153, 154). The best characterized PPIases belong to two families, the cyclophilin-type (Cyp) and the FKB-506 drug-binding protein type (FKBP) (155). Data mining of the *T. brucei* genome identified that Cyp40 and a putative FKB-506 binding like protein (FKBPL) are present in the extracellular parasite proteome (Table 2). Investigation of the domain structure and sequence conservation indicate that both Cyp40 and FKBPL in *T. brucei* were shown to display the characteristic two-domain structure of a N-terminal PPIase domain and a C-terminal TPR domain (Fig 4). Though it must be noted that the C-terminal TPR domain in kinetoplastid Cyp40 underwent substantial evolutionary modification (156), thus potentially impacting Cyp40-Hsp83 interactions. Future structure/function studies should explore the effect these modifications have on the isomerase and chaperone activities of the protein in comparison to its human counterpart.

Studies conducted on the Cyp40 orthologue in *L. donovani* have revealed that the protein functions in *Leishmania* stage-specific morphogenesis, motility, and the development of infectious-stage parasites (156, 157). The study conducted by Yau and colleagues (2014) also suggested that LdCyP40 and LdFKBP2 functions in regulating *Leishmania* cytoskeletal dynamics. Given the capacity of CyP40 and FKBP52 to compete for molecular partners (158), LdCyP40 may interact with microtubules to promote tubulin polymerization as a means of counteracting LdFKBP52-mediated depolymerization. RNAi-mediated knockdown of both Cyp40 and FKBPL in *T. b. brucei* parasites demonstrated that these proteins are essential at the BSF stage and parasite differentiation (58,59,61,85). Proteomic data predicted these proteins to reside in the cytosol and flagellar (70, 135). Together this data indicates that *T. brucei* CyP40 and FKBPL may play essential roles in morphogenesis, motility, and the development of infectious-stage parasites.

#### J52

The J-protein family is a major subset of co-chaperones for the Hsp70 chaperone machinery and they are broadly classified into four subtypes (I-IV). The J-protein family from *T. brucei* has been explored previously (9). It was shown in that study that J52 is one of six type III J proteins in *T brucei* that possesses the TPR domain (others are J42, J51, J52, J53, J65 and J67) (9). J52 is predicted to reside in the cytosol together with J51 and J42 (9). DnaJC7/Tpr2, the human orthologue of J52 was first identified as a cytosolic protein via a two-hybrid screen for interaction with a GAP-related segment (GRD) of neurofibromin. It was reported to encode seven TPR units and possess a domain of high similarity to the DnaJ family (159). Tpr2 also regulates the multichaperone system involving Hsp70 and Hsp90 but in a nucleotide independent manner with Hsp90. DnaJC7 is predominantly thought to be involved in retrograde transport of client proteins from Hsp90 to Hsp70 (160, 161). Proteomic analysis in *T. brucei* showed J52 to be upregulated in the procyclic form of the parasite (58, 59).

### Small glutamine-rich TPR-containing protein (SGT)

The small glutamine-rich TPR-containing protein (SGT) is a co-chaperone involved in a specific branch of the global cellular quality control network that determines the fate of secretory and membrane proteins that mislocalize to the cytosol (162, 163). Human SGT is a modular protein characterized by three characteristic sequence motifs, namely an N-terminal dimerization domain, central TPR domain and a glutamine-rich region at the C terminus (164). The SGT orthologues identified in kinetoplastid protists are atypical (Table 2) as these proteins all lack the characteristic glutamine-rich region and contain a substituted region with charged amino acid residues (165). Proteomic analysis in *T. brucei* identified TbbSGT to be upregulated in the procyclic form of the parasite as well as part of the flagellar and cell surface proteome (58,59,61,70,71). The SGT orthologue in *L. donovani* is an essential protein for *L. donovani* promastigote growth and viability (165). LdSGT was shown to form large, stable complexes that included Hsp83, Hsp70, HIP, HOP, J-proteins, and Hsp100 (165), whereas recombinant *L. braziliensis* SGT was shown to interact with both LbHsp90 and HsHsp70-1A (166). Therefore, the orthologous proteins in *T. b. brucei* and *T. b. gambiense* may have developed the same activity and assist in the formation of the *T. brucei* Hsp83 chaperone system. Though future studies should be conducted to elucidate SGT-Hsp70/Hsp83 interaction in *T. brucei*.

### Non-TPR containing Hsp83 co-chaperones

#### p23

The co-chaperone p23 is a small acidic protein that binds the Hsp90 NBD to stabilise the closed conformation of Hsp90, inhibiting ATPase activity and prevent client protein release from the complex (167, 168). In addition to its HSP90 co-chaperone function, p23 has its own chaperoning activity *in vitro* and can suppress the aggregation of denatured proteins (169, 170). *In silico* analysis of the genomes of both *T. brucei* subspecies revealed that the parasite possesses two evolutionarily divergent p23 orthologues, and subsequently these orthologous proteins were named p23a and p23b (Table 2). The possession of two putative p23 proteins was found to be conserved in all the selected kinetoplastid protists in this study except *B. saltans* (Table 2). The Tbp23a and Tbp23b proteins share 28% identity to each other and share 33% and 26% identity respectively to human p23. Additionally, RNAi knockdown of these proteins showed that each p23 protein is essential to parasite viability at specific stages of the life cycle (Fig 4). The orthologs of these proteins have been explored in two Leishmania *spp.* (171). Both proteins in *L. braziliensis* possessed intrinsic chaperone activity, but they have different client protein specificities; they also inhibit LbrHsp83 ATPase activity to different extents (171). Such functional differences might be important in both Hsp90 regulation and in their interactions with client proteins during the life stage transformations of kinetoplastid parasites. However, to support these assertions, more functional and *in vivo* studies of kinetoplastid p23a and p23b proteins are needed.

#### Aha1

Aha1 has been identified as the primary activator of the ATPase activity of Hsp90 and it acts independent of the other co-chaperones. Homologues of Aha1 have been identified across species from yeast to mammals. Aha1 binds with both its N-and C-terminal domain (Fig 4) to the NBD and MD of Hsp90 to facilitate the dimerization of the chaperone (172–174). Data mining of the *T. brucei* genome identified that the parasite encodes for a single *Aha1* gene (Table 2). The Aha1 orthologue in *L. braziliensis* (LbrAha1) has been characterized, where it was shown to be a cognate protein that shared several structural and functional properties with the human and yeast orthologues. This suggested similar functional mechanisms among these proteins despite the low degree of conservation in the amino acid sequence (131). Recombinant LbrAha1 stimulated the weak ATPase activity of recombinant LbrHsp83 by around 10-fold exhibiting a cooperative behaviour according to the model that two LbrAha1 molecules can act on one LbHsp83 dimer (131). Data from proteomic analysis in *T. brucei* revealed that TbbAha1 is up regulated in the BSF stage of the parasite (59,61,175) as well as being essential to parasite viability at this stage of life cycle (85).

Two other co-chaperones in *T. brucei* had previously been identified, a TPR domain protein identified as Cns1(Tb927.10.11380) and a component of motile flagella 56 (Tb927.9.10490), orthologue of human Pih1 (130). Little has been done to explore these two proteins. So far, only the cytosolic Hsp90 has been shown to require the function of co-chaperones, the other forms of Hsp90 function in the absence of co-chaperones (176, 177).

## Conclusion

The Hsp90 family contains an abundant and essential group of proteins which are highly conserved and implicated in a myriad of cellular functions. Due to their role in cellular proteostasis, they have been implicated in the pathology of many diseases which warrants their targeting as therapeutics (18). Previous studies on the Hsp90 complexes of intracellular kinetoplastids such as *Leishmania* and *T. cruzi* have been conducted (50) but not on the extracellular *T. brucei.* Despite the conservation, distinctive differences exist across species and call for further investigation. In this study we report the *in silico* study of the Hsp90 family and its chaperone complement in *T. brucei*. *T. b. brucei* was found to encode 12 putative Hsp90 proteins, 10 of which are cytosolic (Hsp83). Multiple copies of Hsp83 may allow the parasite to reach a high synthesis level of the proteins in an organism that relies on post-transcriptional regulation and this explains its high levels in the cell even under non-stress conditions (8, 50). The expansion of the Hsp90 chaperone complement also reiterates its importance in the biology and functioning of kinetoplastids (49–51). Hsp83 was also found present in both stages of the parasite but upregulated in the blood stream form (BSF), this is similar to previous findings of much higher transcripts of Hsp83 in blood stream forms of *T. brucei* reflecting their temperature induced role of differentiation (52). The upregulation of Hsp83 together with the co-chaperone Sti1in the BSF may be a further indication of their heat inducibility and involvement in cell defence just as seen in Hsp70 (51).

Hsp90 has been established to partner with co-chaperones to maintain homeostasis, however, Hsp90 seems to partner with the various co-chaperones as dictated by the client being chaperoned (178, 179). This study identified 8 co-chaperones in the *T. brucei* Hsp83 chaperone system which is less than the number of co-chaperones in the human system, confirming that the Hsp90 chaperone machinery is species specific (130). A detailed report for clients in Hsp90 is still largely absent (54). Previous studies have indicated that inhibitors targeting Hsp83 have been shown to cure mice of *T. brucei* infection, although the toxicity of inhibitors to Hsp90 in higher eukaryotes is attributed to a functional loss of client proteins and possible cell cycle arrest (46). Most of the identified Hsp90 client proteins in mammals are kinases (19). Despite the fact that most clients for *T. brucei* Hsp90 have not been identified, over 170 protein kinases (about 30% of the number present in their human host), have been recognised (75, 180). In addition to being regulated by co-chaperones, Hsp90 is also regulated by various post-translational modifications. Some of these PTM sites have been indicated as potential regulatory sites which affect the binding affinity of inhibitors in PfHsp90 (45). The *T. brucei* Hsp90, its co-chaperone network, post-translational modifications, and its regulatory mechanisms as well as the subtle structural differences compared to human Hsp90 all provide a context for a Hsp90-targeted therapy in *T. brucei*.

## Supporting information

**Fig S1. Alignment of the Hsp90/HSPC complement from T. brucei in relation to human and other selected kinetoplastids.**

Multiple sequence alignment of the full-length amino acid sequences was performed using the in-built ClustalW program (81) with default parameters in the MEGA X software (82). Degree of amino acid conservation is symbolized by the following: (*) all fully conserved residues; (:) one of the residues is fully conserved and (.) residues are weakly conserved. The C-terminus motifs are empty-boxed in magenta for the cytosolic Hsp90. Residues involved in post translational modifications accordingly with MS PTM’s proteomic studies by Nett et al, (2009b) and Zhang et al, (2020) coloured red for acetylation and yellow for phosphorylation. The red and yellow empty-boxed sites are highlighting conserved modified residues. Accession numbers for the Hsp90/HSPC amino acid sequences used in this study are provided in Table S1

**Table S1. Accession numbers for the Hsp90/HSPC proteins in *T. brucei* and their respective orthologues in kinetoplastid parasites and *H. sapiens***

*^a^*The accession IDs for the members of the *H. sapiens* Hsp90/HSPC protein family were retrieved from NCBI (https://www.ncbi.nlm.nih.gov/).

*^b^*The accession IDs for the members of the *T. b. brucei* (Tb refers to Tbb), *T. b. gambiense*, *T. cruzi*, *C. fasciculata*, *B. saltans* and *L. major* Hsp90/HSPC protein family were retrieved from the TriTrypDB database (http://tritrypdb.org/tritrypdb/; Aslett et al. 2010).

^c^The accession IDs for the orthologues, identified by reciprocal BLASTP analysis, of three strains of *T. cruzi* are listed. *T. cruzi* CL Brener Esmeraldo-like (TcCLB), *T. cruzi* Dm28c 2018 (C4B63), and *T. cruzi* marinkelli strain B7 (Tc_MARK).

